# Comparison of two injectable anaesthetic protocols in Egyptian fruit bats *(Rousettus aegyptiacus)* undergoing gonadectomy

**DOI:** 10.1101/2021.09.20.461052

**Authors:** Martina Amari, Federica A. Brioschi, Vanessa Rabbogliatti, Federica Di Cesare, Alessandro Pecile, Alessia Giordano, Pierangelo Moretti, William Magnone, Francesco Bonato, Giuliano Ravasio

## Abstract

Egyptian fruit bats are experimental animals of increasing interest because they have been identified as a natural reservoir for several emerging zoonotic viruses. For this reason, bats could undergo different experimental procedures that require sedation or anaesthesia. Our aim was to compare the effects of two balanced anaesthetic protocols on sedation, cardiopulmonary variables and recovery in bats undergoing gonadectomy. Twenty bats were randomized into two groups; patients in group DK received intramuscular injection of dexmedetomidine (40 μg kg^-1^) and ketamine (7 mg kg^-1^), whereas those in group DBM were anaesthetized with intramuscular dexmedetomidine (40 μg kg^-1^), butorphanol (0.3 mg kg^-1^) and midazolam (0.3 mg kg^-1^). Time of induction, cardiopulmonary parameters and anaesthetic depth were measured. If anaesthesia plan was considered inadequate, fraction of inspired isoflurane was titrate-to-effect to achieve immobility. At the end of the surgery venous blood gas analysis was performed and intramuscular atipamezole (200 μg kg^-1^) or atipamezole (200 μg kg^-1^) and flumazenil (0.03 mg kg^-1^) was administered for timed and scored recovery phase. A significantly higher heart rate and peripheral oxygen saturation were recorded in DBM group (*p* = 0.001; *p* = 0.003 respectively), while respiratory rate was significantly lower than DK group (*p* = 0.001). All bats required isoflurane supplementation during surgery with no significant difference. No differences were observed in rectal temperature, induction and recovery times. Sodium and chlorine where significantly higher in DBM group (*p* = 0.001; *p* = 0.002 respectively). Recovery scores in group DK were significantly better than in group DBM (*p* = 0.034). Both protocols induced anaesthesia in Egyptian fruit bats with comparable sedative and cardiorespiratory effect. These drug combinations may be useful for minor procedures in bats, and they could be associated with inhalation anaesthesia in determining and maintaining a surgical anaesthetic plan.

## Introduction

Pteropid bats have been studied in various research fields as they have been identified as a natural reservoir for various emerging zoonotic viruses, including Marburg virus [1], Hendra virus, Nipah virus [2] and lyssavirus variants [3]. Moreover, among the family *Pteropodidae*, also the Egyptian fruit bat (*Rousettus aegyptiacus*) showed characteristics of a reservoir host for SARS-CoV-2 [4]. Besides, Egyptian fruit bats are commonly housed in zoological environments because they are small, amenable to handling and reproduce readily in captivity [5].

The safe collection of biological samples from pteropid bats, such as blood and swabs from the throat, urethra and rectum, is essential for both the animal and the operator [6]. During sampling, the physical restraint of bats can expose the handler to bite and scratch injuries, resulting in potential zoonoses transmission. Short-term anaesthesia facilitates operator safety and minimises stress for the bat [7,8]. Moreover, in zoological settings, short-term anaesthesia results important to apply contraception protocols, to prevent overpopulation and inbreeding in highly fertile bat colonies [9,10].

A total isoflurane inhalation anaesthesia is often the method of choice for bats, having the advantage of wide safety margin, very little metabolization, and quick induction and recovery times [11,12]. However, isoflurane is not commonly used under field conditions [13] and it does not provide a sufficient analgesic action [14]. Side effects that may occur with extensive inhaled anaesthetic use are dose-dependent and include respiratory and myocardial depression and decreased in sympathetic activity, leading to decreased cardiac output and hypotension [15]. For this reason, halogenates are often combined with injectable anaesthetics to reduce anaesthetic requirements and cardiopulmonary effects [16,17].

Previously reported injectable anaesthetic protocols used in bats include alpha-2 adrenergic agonists (i.e. xylazine or medetomidine) and ketamine (KET) [6–8,18]. The use of alpha-2 adrenergic agonists results in sedation, analgesia, muscle relaxation, and anxiolysis, and reduces the anaesthetic requirements of injectable and inhalant agents during induction and maintenance of general anaesthesia [19]. Dexmedetomidine (DEX) is the alpha-2 agonist with the highest receptor selectivity and it is twice as potent as medetomidine [20].

Ketamine induces anaesthesia and amnesia by functional dissociation of the central nervous system resulting in catalepsy, immobility, amnesia, and marked analgesia [19], but its use alone is highly discouraged due to the poor muscle relaxation and slow and often excitative awakenings [18]. The alpha-2 adrenergic agonists-KET combination provide good analgesia and muscle relaxation together with an excellent cardiovascular stability, but it may be associated with prolonged recovery and hypothermia [21].

Butorphanol (BUT) is a k agonist-μ antagonist opioid with mild sedative and analgesic properties [19]. Opioids are often combined with alpha-2 adrenergic agonists because they potentiate their sedative and analgesic effects with minimal additional cardiovascular effects [22].

Midazolam (MDZ) is a benzodiazepine that has sedative-hypnotic, anxiolytic and muscle relaxant effects and enhances the sedative and antinociceptive effects of alpha-2-adrenergic agonists [23].

A subcutaneous combination of medetomidine, MDZ and opioids has been shown to be safe for Egyptian fruit bat anaesthesia, with no apparent morbidity or mortality [21].

The purpose of the study was to evaluate and compare sedative effects of two different injectable anaesthetic protocols, DEX and KET (group DK) versus DEX, BUT and MDZ (group DBM) in bats undergoing gonadectomy and to record physiological and adverse effects following administration of both protocols. We also evaluated the duration of induction and timing and quality of recovery achieved by both combinations. We hypothesized that both injectable protocols are effective in determining and maintaining a surgical anesthetic plan with no or diminished use of isoflurane in Egyptian fruit bats with few side effects.

## Materials and methods

### Ethics statement

The present study complies with ethical standards and was conducted under the approval of the Institutional Ethical Committee for Animal Care at the University of Milan (OPBA_104_2021). Owner’s written informed consent was obtained.

### Animals and housing conditions

Twenty healthy male and female Egyptian fruit bats (age unknown, body weight between 100 and 150 g) presented at the Veterinary Teaching Hospital of the University of Milan to perform gonadectomy were included in the study.

All bats were housed together throughout the hospitalization period in a large mesh cage (height 2.5 m, width 1.5 m, length 2.0 m) in a controlled-environment room (22-25 °C and 60% humidity) and exposed to a natural photoperiod (light/dark alternation period of 8/16 hours). They received water *ad libitum* and were fed with a mixture of seasonal fruits and vegetables. All the procedures were performed after an acclimation period of 10 days. During this period, they were considered healthy based on observation of normal behaviour without stereotypic attitudes, normal activity levels and appetite along with normal weight, size and wingspan length, in absence of clinical signs.

### Study design

The day of the surgery each bat was captured inside the cage using protective leather gloves and was temporarily placed inside a perforated canvas bag on a digital laboratory scale (Precisa BJ610C, Precisa Instrument, Dietikon, Switzerland) to accurately measure its body weight. Patients were then randomly assigned to either group DK or DBM (www.randomizer.org). Bats in group DK received an intramuscular (IM) administration of DEX (40 μg kg^-1^)(Dexdomitor 0.5 mg ml^-1^; Vetoquinol Italia S.r.l., Italy) and KET (7 mg kg^-1^) (Lobotor 100 mg ml^-1^; ACME S.r.l., Italy) and those in group DBM received an IM injection of DEX (40 μg kg^-1^), BUT (0.3 mg kg^-1^)(Nargesic 10 mg ml^-1^; ACME S.r.l., Italy) and MDZ (0.3 mg kg^-1^)(Midazolam Hameln 5 mg ml^-1^; Hameln pharma gmbh, Germany). All syringes were prepared, labelled in a way that did not reveal their content and injected in the bats’ thigh muscles by an experienced anaesthetist not involved in the study. All anaesthetic procedures were performed by another experienced anaesthetist, who was unaware of the treatment administered.

Immediately after drugs’ administration, the bat was placed in a transparent plexiglass cage and observed continuously to monitor the induction phase and record the induction time. The induction time was defined as the interval from administration of the drugs to the absence of movement following a gentle foot palpation.

Upon the loss of response following palpation, the bat was positioned in dorsal recumbency on a warm air blanket (Bair Hugger 505 Warming Unit; 3M, Germany), which was covered with an adsorbent pet sheet. Moreover, a monopolar electrosurgical plate was placed under the bat. A complete physical examination was performed, and the wingspan and the body length (head to tail) were measured. Based on the size of the animal and development of the reproductive system, the age of each bat was estimated and classified into “juvenile” or “adult”. Adult males were distinguished on the basis of fully developed penis and testes, a body size ≥ 15 cm and a wingspan ≥ 48 cm; adult females were distinguished from juvenile on the basis of worn or enlarged nipples or if it were palpably pregnant, a body size ≥ 14 cm and a wingspan ≥ 48 cm. Juveniles (< 12 months old) were classified on their smaller size and rudimentary development of sexual characteristics [24]. A 22-gauge venous catheter (Jelco IV Catheter Radiopaque; Smiths Medical Italia S.r.l., Italy) was inserted in the left cephalic vein. A multiparameter monitor (S5 Compact Anesthesia Monitor; Datex-Ohmeda, USA) was used throughout the anaesthetic period. Oxygen 100% flow-by at 1 L min^-1^ was administered via a facemask, which was attached to a side-stream spirometer. A pulse oximeter was connected on the right hind leg and disposable foam pad electrodes for electrocardiographic measurements were positioned as in Fig 1.

**Fig 1.**
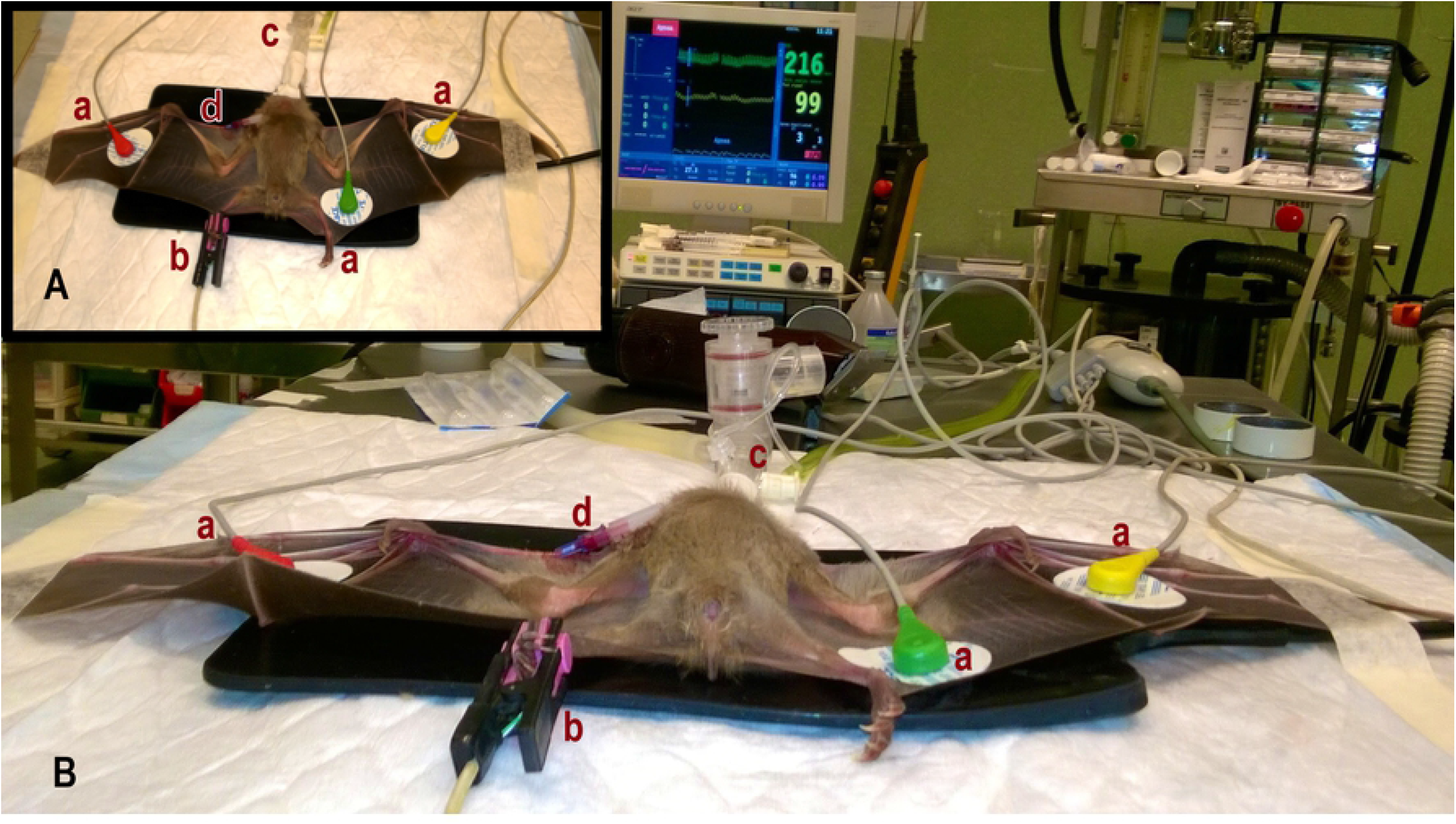
Egyptian fruit bat in dorsal recumbency, instrumented with monitoring devices (A) and overview in operating room (B). Disposable foam pad electrodes (a) positioned on the ventral aspect of wings for electrocardiographic measurements; pulse oximetry probe (b) placed on the right hind leg for haemoglobin saturation measurement; side-stream spirometer (c) attached to the facemask for multi gas analysis, and 22-gauge catheter (d) inserted in the left cephalic vein for collection of samples for blood gas analysis.

Heart rate (HR), respiratory rate (RR) and peripheral oxygen saturation (SpO_2_) were continuously monitored and recorded every 5 minutes during surgery. Rectal temperature (RT) was measured at the beginning and at the end of the surgical procedure using a digital thermometer (Pic VedoFamily; Pikdare S.p.A., Italy).

Depth of anaesthesia were assessed every 10 minutes by easy extension and flexion of the wing without any voluntary movement or presence of muscle tone by opening the jaw. In case of spontaneous movement or presence of muscle tone, the anaesthesia depth was considered inadequate, and the fraction of inspired isoflurane (FI-ISO) was titrate-to-effect to the minimum concentration to achieve immobility and loss of muscle tone and this value was recorded and then adjusted over time as needed.

All gonadectomy surgeries were performed by the same experienced surgeon and total surgery time was recorded. Females that were found to be pregnant underwent ovariohysterectomy.

During the entire procedure, side effects including arrhythmia, irregular breathing pattern, twitching and tremors were recorded as existing or not, regardless of severity or duration.

At the end of the surgery, venous blood gas analysis (Stat Profile pHOx Ultra; Nova biomedical Italia S.r.l., Italy) was performed. Analysis included venous pH, venous partial pressure of oxygen (PvO_2_) and carbon dioxide (PvCO_2_), base excess (BE) and electrolytes (Na^+^, K^+^, Cl^-^) as well as bicarbonate (HCO^3-^) and total haemoglobin (Hb). Then, bats in group DK received IM atipamezole (200 μg kg^-1^) (Antisedan 5 mg ml^-1^; Vetoquinol Italia S.r.l., Italy), while bats in group DBM received IM atipamezole (200 μg kg^-1^) and IM flumazenil (0.03 mg kg^-1^) (Flumazenil Kabi 0.1 mg ml^-1^; Fresenius Kabi Italia S.r.l., Italy). Following the IM injection of reversal drugs in the thigh muscles, each bat was returned in the plexiglass cage.

Recovery time, namely the time from the injection of the antagonists to flying, was recorded and recovery quality scored on a scale of 1-3 (Table 1). All the recoveries were observed continuously and evaluated by the same anaesthetist.

**Table 1.**
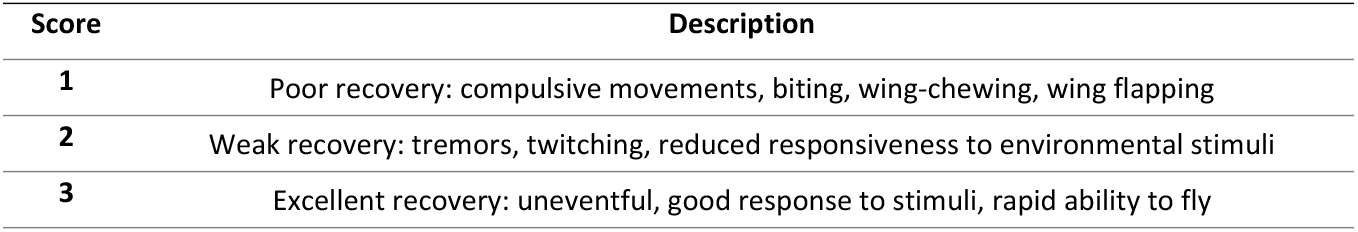
Scoring system used to assess recovery from anaesthesia.

| Score | Description |
| --- | --- |
| **1** | Poor recovery: compulsive movements, biting, wing-chewing, wing flapping |
| **2** | Weak recovery: tremors, twitching, reduced responsiveness to environmental stimuli |
| **3** | Excellent recovery: uneventful, good response to stimuli, rapid ability to fly |

After recovery, all bats were monitored every hour until 12 hours, and then were observed daily for a week to evaluate any side effects.

### Statistical analysis

A power analysis was performed and determined that a minimum of 18 bats would be required to detect a clinically relevant difference in induction time of 4 minutes or more between the two groups with a power of 85% and α = 0.05 (two-tailed).

Statistical analysis was performed using IBM SPSS Statistics 26.0 (SPSS Inc, Chicago, USA). The normality of data distribution was assessed by a Shapiro-Wilk test at the α = 0.05 level. Descriptive statistics were reported as mean ± standard deviation (SD) or median (range) for continuous and ordinal variables, respectively. Pearson’s chi-squared test was used to evaluate significant differences in nominal data. Analysis of variance (ANOVAs), followed by Bonferroni’s post hoc test, and Mann-Whitney U test or Wilcoxon’s test was applied for normal and non-normal data, respectively, to assess significant differences between and within groups. The influence of total surgery time on recovery time was evaluated by Pearson’s correlation. Differences with *p* < 0.05 were considered significant.

## Results

Twenty Egyptian fruit bats were included in the study: ten bats (5 males, 5 females) received DK treatment and ten bats (4 males, 6 females) received DBM treatment. No significant differences in gender, age (DK 8 adults, 2 juveniles; DBM 7 adults, 3 juveniles), female reproductive status (DK 5 pregnant; DBM 5 pregnant) and body weight (DK 111.4 ± 9.38 g; DBM 111.6 ± 6.91 g) were recorded. There were no significant differences in mean induction time and in total surgery time between the two treatment groups (Table 2).

**Table 2.** Induction time, surgery time and recovery time in 20 Egyptian fruit bats anaesthetized for gonadectomy.

|  | DK | DBM |
| --- | --- | --- |
| **Induction time (seconds)** | 149 ± 170 | 169 ± 159 |
| **Surgery time (minutes)** | 53 ± 16 | 44 ± 12 |
| **Recovery time (seconds)** | 345 ± 150 | 424 ± 210 |

Bats in group DK ( $n = 10$ ) received DEX and KET combination and bats in group DBM ( $n = 10$ ) received DEX, BUT and MDZ administration. Atipamezole (group DK) or atipamezole and flumazenil combination (group DBM) was administered intramuscularly at the end of the surgery.
Results are presented as mean ± standard deviation.

Heart rate, RR and SpO_2_ were compared between groups for the first 50 minutes following induction (on further time-points some of the bats had already recovered). A significantly higher heart rate was recorded in DBM group (DK 181 ± 31 bpm; DBM 203 ± 47 bpm) (*p* = 0.001), while respiratory rate was significantly lower than DK group (DK 112 ± 26 bpm; DBM 85 ± 21 rpm) (*p* = 0.001). A significant difference was observed in peripheral oxygen saturation, where in the DBM group it was higher than in the DK group (DK 98.1 ± 1.9 %; DBM 99.1 ± 0.9 %) (*p* = 0.003). All bats required isoflurane supplementation during surgery and no significant difference in FI-ISO was observed between groups. No statistically significant differences were observed within groups over time in HR, RR, SpO_2_ and FI-ISO parameters. Results are summarized in Figs 2 and 3.

**Fig 2.**
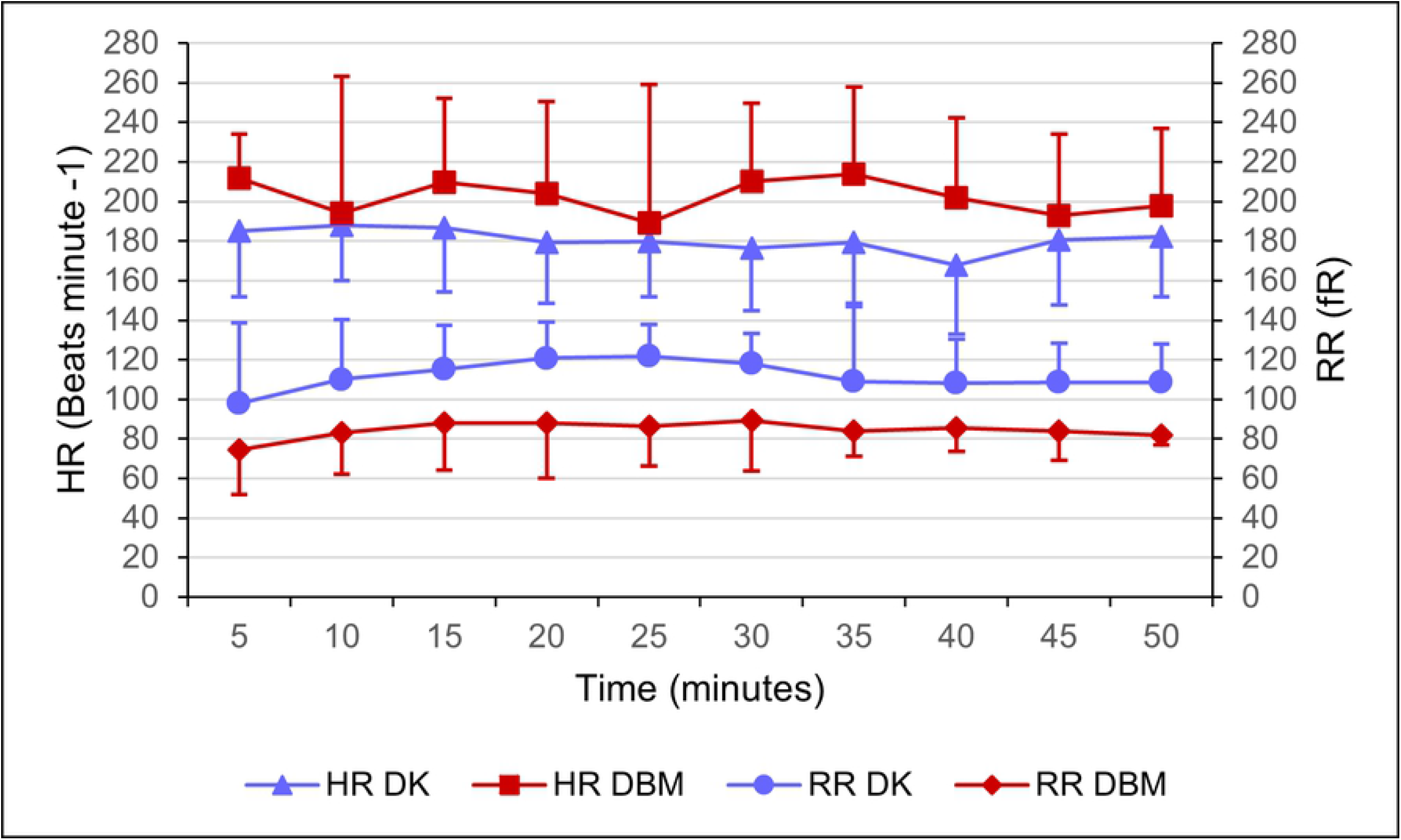
Heart rate (HR) and respiratory rate (RR) in 20 Egyptian fruit bats during general anaesthesia for gonadectomy. Bats in group DK (*n* = 10) received DEX and KET combination and bats in group DBM (*n* = 10) received DEX, BUT and MDZ administration. Results are presented as mean ± standard deviation. Significant differences (*p* < 0.05) between groups in HR and RR were found.

**Fig 3.**
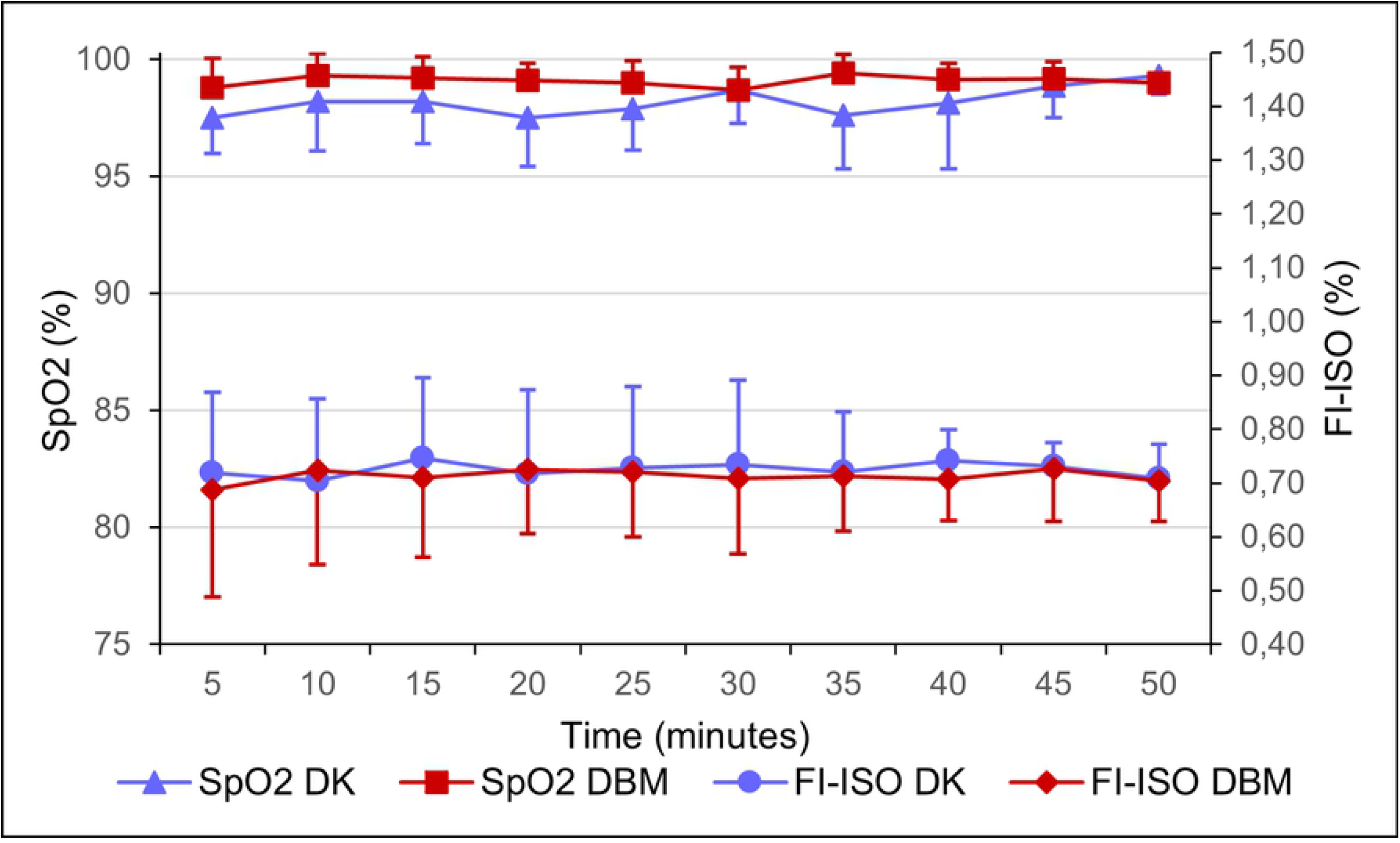
Peripheral oxygen saturation (SpO_2_) and fraction of inspired isoflurane (FI-ISO) in 20 Egyptian fruit bats during general anaesthesia for gonadectomy. Bats in group DK (*n* = 10) received DEX and KET combination and bats in group DBM (*n* = 10) received DEX, BUT and MDZ administration. Results are presented as mean ± standard deviation. Significant differences (*p* < 0.05) between groups in SpO_2_ were found.

There were no significant differences in initial (DK 37.5 °C ± 0.7; DBM 37.7 °C ± 0.7) or final (DK 36.8 °C ± 1.3; DBM 37.2 °C ± 1.1) rectal temperature between treatments and RT at the end of the surgery did not decrease significantly compared to the beginning of the surgical procedures in either group.

There was no significant difference between groups in venous blood gas analysis except for Na^+^ (mmol L^-1^) (*p* = 0.001) and Cl^-^ (mmol L^-1^) (*p* = 0.002) that where significantly higher in DBM group (Table 3).

**Table 3.** Venous blood-gas values, venous pH, total haemoglobin, venous base excess, venous bicarbonate, and electrolytes in 20 Egyptian fruit bats anaesthetized for gonadectomy.

| Parameter | Group | Mean ± SD |
| --- | --- | --- |
| **PvCO<sub>2</sub> (mmHg)** | DK | 39.10 ± 6.4 |
| | DBM | 40.90 ± 5.0 |
| **PvO<sub>2</sub> (mmHg)** | DK | 237.80 ± 137.0 |
| | DBM | 277.00 ± 117.1 |
| **pH** | DK | 7.35 ± 0.04 |
| | DBM | 7.35 ± 0.02 |
| **Hb (g dL<sup>-1</sup>)** | DK | 13.70 ± 0.88 |
| | DBM | 13.48 ± 1.33 |
| **BE (mmol L<sup>-1</sup>)** | DK | -4.10 ± 1.47 |
| | DBM | -3.06 ± 2.05 |
| **HCO<sub>3</sub><sup>-</sup> (mmol L<sup>-1</sup>)** | DK | 20.73 ± 1.00 |
| | DBM | 21.81 ± 1.59 |
| **Na<sup>+</sup> (mmol L<sup>-1</sup>)** | DK | 134.8 ± 1.75* |
| | DBM | 139.5 ± 1.58* |
| **K<sup>+</sup> (mmol L<sup>-1</sup>)** | DK | 3.67 ± 0.52 |
| | DBM | 3.65 ± 0.66 |
| **Cl<sup>-</sup> (mmol L<sup>-1</sup>)** | DK | 103.40 ± 1.78* |
| | DBM | 107.50 ± 2.42* |

Bats in group DK (*n* = 10) received DEX and KET combination and bats in group DBM (*n* = 10) received DEX, BUT and MDZ administration. Results are presented as mean ± standard deviation (SD).
PvCO<sub>2</sub>, PvO<sub>2</sub>, venous oxygen and carbon dioxide partial pressures; Hb, total haemoglobin; BE, venous base excess; HCO<sub>3</sub><sup>-</sup> venous bicarbonate; Na<sup>+</sup> ionized sodium; K<sup>+</sup> ionized potassium; Cl<sup>-</sup> ionized chlorine.
Significant differences (*p* < 0.05) between groups are indicated by \*.

Recovery times did not significantly differ between groups and no correlation was observed between total surgery time and recovery duration (Table 2).

A significantly worse recovery quality was observed in the DBM group (DK median 3, range 3-3 and DBM median 3, range 2-3) (*p* = 0.034) as shown in Fig 4.

**Fig 4.**
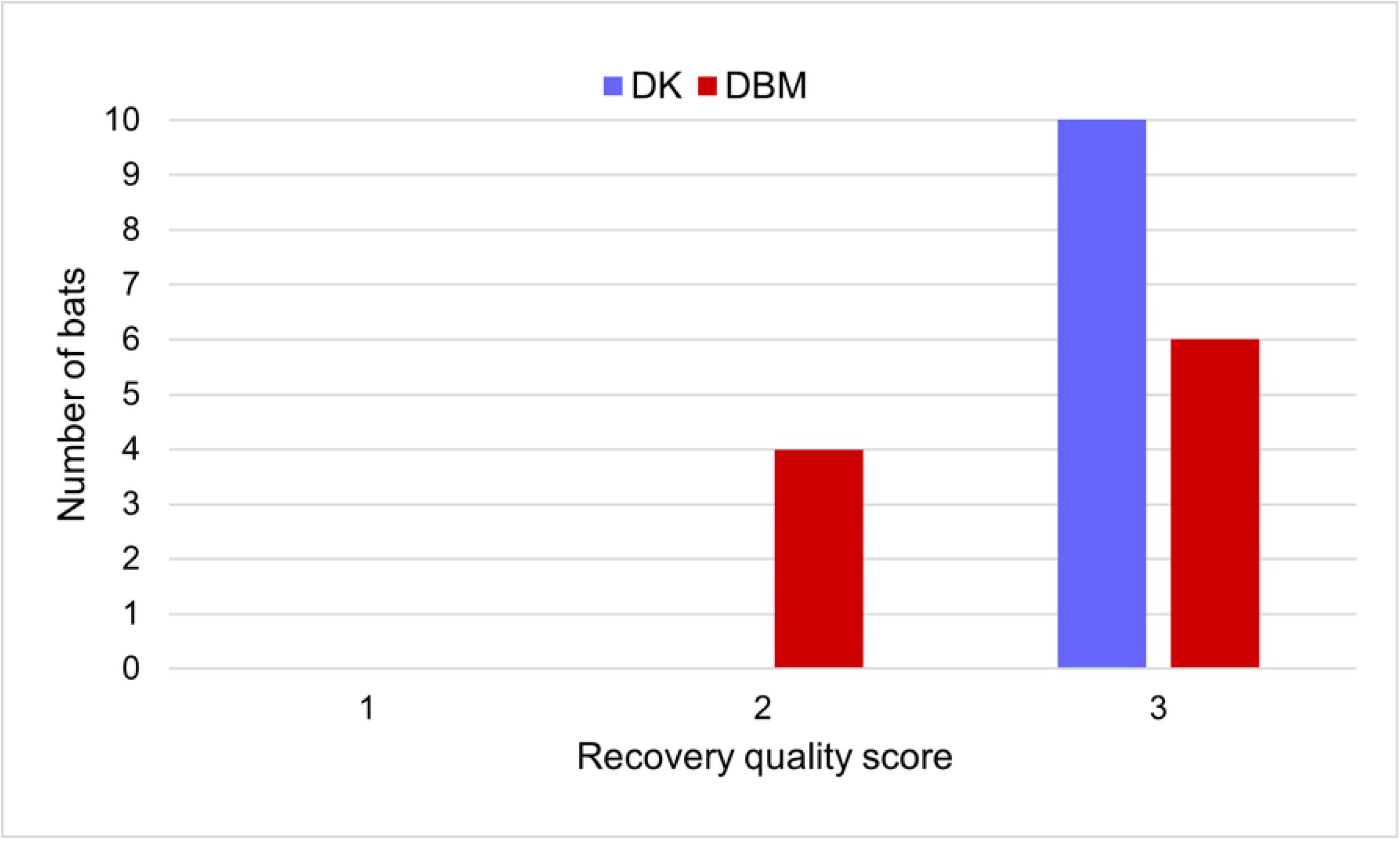
Recovery quality scores in 20 Egyptian fruit bats anaesthetized for gonadectomy. Bats in group DK (*n* = 10) received DEX and KET combination and bats in group DBM (*n* = 10) received DEX, BUT and MDZ administration. Significant differences (*p* < 0.05) between groups were found.

No side effects, such as arrhythmia or irregular breathing pattern, twitching and tremors, were observed in any bat following intramuscular administration and during the surgery. After recovery, all bats returned to normal behaviour and good activity levels and appetite; no side effects were observed during the follow-up period.

## Discussion

The present study aimed to evaluate sedative effects of two different injectable anaesthetic protocols, DEX and KET (DK) versus DEX, BUT and MDZ (DBM) in bats undergoing gonadectomy.

Different studies report the use of alpha-2 adrenergic agonists in association with KET in bats [6–8,18]. Dexmedetomidine-ketamine combinations have been used in a variety of mammalian species [25–27]. To the authors’ knowledge, no studies have been carried out in Egyptian fruit bats, or any other Chiroptera species, using DEX as a part of a balanced anaesthetic protocol. In the present study, this association produced high-quality immobilization with rapid and smooth induction.

Anaesthesiologic protocols including the association of alpha-2 adrenergic agonists, opioids and benzodiazepines have already been described in veterinary medicine in several species [28,29]. As regards bats, only Tuval and colleagues (2018) have compared different subcutaneous combinations of medetomidine, MDZ and opioids in *R. aegyptiacus* with no apparent morbidity and mortality. Compared with the work of Tuval and colleagues (2018), the dosages and total volumes of drugs used in this study were considerably lower, probably also due to the different route of administration. In our study, IM administration was performed in the thigh muscles; no behaviour referable to muscle soreness or pain was observed following injection or during recovery phase, and no bats had difficulty in flying following the procedure.

A rapid and gentle induction was observed in all bats in the DBM group, and they reached the desired level of sedation in slightly longer time than the DK group, but with less intragroup variability. The association of these three drug classes, at lower doses than would be made if only one agent were used, results in a synergistic central nervous system depressant response while minimizing the undesirable side effects of each drug [30]. Further advantage of this association is that each drug can be completely antagonized enabling precise timing of sedative effects and in case of emergency condition, allowing for safer patient management. In the present study, butorphanol reversal was not performed to preserve post-operative pain management in all bats included in DBM group.

All bats required isoflurane supplementation without differences in FI-ISO between groups and it only became necessary during the surgical phase, while injectable anaesthetics were satisfactory for the patient preparation. The use of 2-2.5% isoflurane via facemask for the maintenance of general anaesthesia in Chiroptera species has been described in several works [9,11–13]. In the present study, the FI-ISO necessary to obtain an adequate surgical plane of anaesthesia was much lower than that reported in literature, suggesting that both protocols may have had a sparing effect on isoflurane. The minimal alveolar concentration reduction of inhaled anaesthetics after administration of DEX, KET, BUT and MDZ is reported in various species [20,31,32]. The results of this study suggested that to obtain an adequate plane for surgical anaesthesia during gonadectomy, isoflurane had to be administered, even if at lower concentrations than those reported in the literature. However, in the authors’ opinion, for minor procedures (physical examination, manipulation, blood and swab sampling or skin biopsy) both these injectable anaesthesia protocols could be sufficient to achieve immobility in bats without stress and were used safely by the operator during the entire anaesthetic period.

Noll and colleagues (1979) reported a resting HR of 248 ± 3 bpm in telemetrically monitored adults of *R. aegyptiacus* [33]. In the present study, baseline values in manually restrained bats before drugs administration were not measured. However, physical restraint is stressful and would have altered values themselves, as reported in previous studies [34]. Therefore, considering the resting parameters in the literature, it is possible to assume that there was a decrease in HR following the administration of both injectable protocols and this finding is probably imputable to the effect of DEX. The cardiac effects of DEX observed in this study are the same as those described for other mammalian species and other alpha-2 agonists [20,21,25]. Dose-dependent bradycardia following DEX administration results primarily from a decrease in sympathetic tone and partly by baroreceptor reflex and enhanced vagal activity [20]. Opioids decrease HR by increasing parasympathetic tone [19], while KET should balance the effects on the cardiovascular system induced by alpha-2 adrenergic agonists [25]. Nevertheless, the use of adjunctive drugs, such as alpha-2 adrenergic agonists, tends to blunt the sympathomimetic effect of KET and to decrease cardiac function and arterial blood pressure [19]. Indeed, in this study, HR was significantly lower in DK group than DBM group.

Comparing baseline values of RR reported by Tuval and colleagues (2018), in the present study both protocols showed a slight reduction in the values of this physiological parameter if compared to other studies [21,34]. Bats in group DBM showed the greatest reduction in RR during anaesthesia, being significantly lower than which occurred in group DK. However, SpO_2_ was significantly higher in the DBM group, although in both groups the measured values were within the normal ranges. Oxygen saturation values have always remained within normal ranges probably due to the reduction in isoflurane concentration [32] and because 100% oxygen was administered throughout the anaesthesia period, as oxygen supply was reported to improve arterial oxygenation during anaesthesia in other species [35]. In the present study, arterial blood gas analysis could not be performed, but peripheral venous blood drawn anaerobically is known to correlate reasonably well with arterial values, at least for pH, bicarbonate and carbon dioxide tension values (CO_2_) [36]. Kelly et al. (2005) showed that a PvCO_2_ of less than 45 mmHg has a 100% negative predictive value to rule out arterial partial pressure of CO_2_ (PaCO_2_) greater than 50 mmHg in humans [37]. Therefore, normal peripheral PvCO_2_ can be used as a screen to exclude hypercapnic respiratory disease [38] and in the present study, hypoventilation induced by inhaled and injectable anaesthetics was neither reflected in changes in PvCO_2_, pH, or PvO_2_ outside physiological ranges in other mammalian species [39] nor in significant differences between the two groups.

Blood gas analysis did not show significant differences in the other values, except for electrolytes, where Na^+^ and Cl^-^ were significantly higher in DBM group. However, no references have been found in the literature to explain a correlation between the plasma concentration of these ions and the effects induced by anaesthetic drugs. So, in healthy animals this difference could be reasonably attributed to animal feeding and or to water intake [40].

Due to the anatomical conformation of the wings and the small size with an high ratio between surface and body mass, Chiroptera are particularly susceptible to heat loss [13,21]. Rectal temperature was measured at the beginning and at the end of the surgical procedure and all bats were warmed with heating pad during general anaesthesia. No significant differences between initial and final temperatures within and between groups were found, suggesting that active warming counteracted the hypothermia induced by general anaesthesia.

In both groups, the simultaneous use of several drugs, with their synergistic effects, may have contribute to achieve an excellent and rapid anaesthetic plan, to reduce the single drug dosages and to avoid the appearance of side effects, as reported by other authors [30,41]. Indeed, no complications associated with the balanced anaesthesia have been observed in the two groups during induction, maintenance, and recovery from general anaesthesia.

Recovery time never exceeded 11 minutes, without significant differences between groups, which suggests that the use of antagonists (atipamezole and atipamezole/flumazenil) at the end of the surgery is advisable to ensure a rapid recovery. In 4 out of 10 bats of DBM group tremors and twitching were observed during recovery along with a greater difficulty in recovering normal wakefulness and responsiveness to environmental stimuli, probably due to the residual sedative effect of BUT. This result is similar to what observed by Tuval *et al*. (2018), where the recovery of bats administered a combination of medetomidine, MDZ and BUT were significantly longer than other groups. Therefore, they concluded that the administration of atipamezole following the use of a protocol containing an alpha-2 adrenergic agonist and BUT in Egyptian fruit bats is recommended.

A limitation of the present study is that the depth of anaesthesia was assessed without evaluating reflexes and probably prevented us from detecting small differences in isoflurane requirements between groups. In addition, although FI-ISO was accurately recorded, no bat was intubated, and it was not possible to determine intraoperative end-tidal concentrations of isoflurane. Further studies are justified to evaluate minimal alveolar concentration reduction induced by both protocols. Finally, we did not evaluate and compare the quality of analgesia induced by both protocols, although changes in cardiorespiratory parameters possibly related to nociception were never recorded in any bat during surgery.

## Conclusions

In conclusion, DK and DBM protocols induced anaesthesia in Egyptian fruit bats with comparable sedative and cardiorespiratory effects and without apparent morbidity or mortality. As Egyptian fruit bats are of increasing interest as experimental animals due to their role as virus reservoirs, chemical restraint of this species is becoming increasingly important to improve research in this field. These drug combinations may be useful for minor procedures in Egyptian fruit bats, and they could be associated with inhalation anaesthesia in determining and maintaining a surgical anaesthetic plan.

## Acknowledgments

We thank the staff of “The Giant” involved in the supply of fresh fruit for patient nutrition and students for their precious and enthusiastic contribution in bats care.

